# Mayaro Virus Infection Elicits an Innate Immune Response in *Anopheles stephensi*

**DOI:** 10.1101/2020.11.15.383596

**Authors:** Cory Henderson, Marco Brustolin, Shivanand Hegde, Gargi Dayama, Nelson Lau, Grant L. Hughes, Christina Bergey, Jason L. Rasgon

## Abstract

Mayaro virus (MAYV) is an arboviral pathogen in the genus *Alphavirus* that is circulating in South America with potential to spread to naïve regions. MAYV is also one of the few viruses with the ability to be transmitted by mosquitoes in the genus *Anopheles*, as well as the typical arboviral transmitting mosquitoes in the genus *Aedes*. Few studies have investigated the infection response of *Anopheles* mosquitoes. In this study we detail the transcriptomic and small RNA responses of *An. stephensi* to infection with MAYV via infectious bloodmeal at 2, 7, and 14 days post infection (dpi). 487 unique transcripts were significantly regulated, 78 putative novel miRNAs were identified, and an siRNA response is observed targeting the MAYV genome. Gene ontology analysis of transcripts regulated at each timepoint suggested activation of the Toll pathway at 7 dpi and repression of pathways related to autophagy and apoptosis at 14 dpi. These findings provide a basic understanding of the infection response of *An. stephensi* to MAYV and help to identify host factors which might be useful to target to inhibit viral replication in *Anopheles* mosquitoes.

**AUTHOR SUMMARY:** Mayaro virus (MAYV) is a mosquito-borne Alphavirus responsible for outbreaks in South America and the Caribbean. In this study we infected *Anopheles stephensi* with MAYV and sequenced mRNA and small RNA to understand how MAYV infection impacts gene transcription and the expression of small RNAs in the mosquito vector. Genes involved with innate immunity and signaling pathways related to cell death are regulated in response to MAYV infection of *An. stephensi*, we also discover novel miRNAs and describe the expression patterns of miRNAs, siRNAs, and piRNAs following bloodmeal ingestion. These results suggest that MAYV does induce a molecular response to infection in its mosquito vector species.

## INTRODUCTION

Mayaro virus (MAYV) is a mosquito-borne, enveloped positive-sense single-stranded RNA virus in the genus *Alphavirus*, first isolated from the blood of five febrile workers in Mayaro county, Trinidad in 1954 [1]. Symptoms of MAYV infection are similar to other arboviral infections such as Dengue (DENV) or Chikungunya viruses, (CHIKV) and include rash, fever, retro-orbital pain, headache, diarrhea, and arthralgia [2]. While no epidemics or outbreaks with MAYV being the causative agent have been recorded outside of South America, there have been imported cases reported in the Netherlands, Germany, France, and Switzerland [3–6], which demonstrates a need to understand the capacity for the virus to spread into naïve regions, such as the United States.

The principal mosquitoes transmitting MAYV naturally are thought to be the canopy-dwellers of the genus *Haemogogus*, maintaining the sylvatic cycle between non-human primates as primary hosts and birds as secondary hosts [7]. Human infections are sporadic due to the rare display of anthropophilic biting behaviors by *Haemogogus* mosquitoes, with transmission due to these species primarily occurring in rural regions with close proximity to forests [8]. Vector competence studies have identified anthropophilic and urban adapted species such as *Aedes aegypti* and *Ae. albopictus*, as well as the malaria parasite transmitters *Anopheles gambiae, An. stephensi, An. freeborni*, and *An. quadrimaculatus*, as being competent vectors for MAYV under laboratory conditions [9–12]. Transmission of an arbovirus with an anopheline mosquito as a primary vector is rare, only having been observed occurring regularly for O’nyong’nyong virus by *An. gambiae* and *An. funestus* in Uganda [13], with some limited evidence for CHIKV and Semliki Forest virus (SFV) [14].

As arboviral pathogens are transmitted between hosts primarily by arthropod vectors, transmission requires the virus to infect and disseminate from the midgut and salivary glands of the mosquito following an infectious bloodmeal [15]. The molecular underpinnings controlling why MAYV and these closely related viruses can infect *Anopheles* salivary glands is of epidemiological interest, yet remains poorly understood. A more complete understanding of this phenomenon requires investigation of the molecular pathways involved in viral infection of anopheline mosquitoes. Recent transcriptomic studies have identified a number of genes involved in classical immune pathways, RNA interference (RNAi), metabolism, energy production, and transport as being regulated in response to arboviral infection of mosquitoes [16–19]. In addition, studies focusing on small RNA identification and regulation have identified RNAi activity, such as miRNA, piRNA, and siRNA expression, in response to infection of mosquitoes by arboviruses [20–23].

The available evidence suggests that, should MAYV be introduced into a naïve region, outbreaks and epidemics of the resulting disease could be driven by anopheline vectors [9, 24–25]. Because anopheline, and not aedine, mosquitoes could act as the primary transmitting vectors for MAYV, this study also provides an opportunity to understand how vector competence might emerge in this system and provide insight into why *Anopheles* are generally poor viral transmitters when compared to *Aedes* mosquitoes. We used RNA sequencing to study the transcriptomic and small RNA responses of *An. stephensi* to infection with MAYV via infectious bloodmeal at 2, 7, and 14 days post infection (dpi).

## MATERIALS AND METHODS

### *Anopheles stephensi* Rearing

Protocols pertaining to mosquito rearing and presentation of infectious bloodmeal has been described elsewhere [9]. Briefly, *An. stephensi* (Liston strain) were provided by Johns Hopkins University (Baltimore, MD, USA). Mosquito colonies were reared and maintained at the Millennium Sciences Complex insectary (The Pennsylvania State University, University Park, PA, USA) at 27°C ± 1°C, 12 hour light 12 hour dark diurnal cycle at 80% relative humidity in 30×30×30-cm cages. Ground fish flakes (TetraMin, Melle, Germany) were used to feed larvae, and upon emergence adult mosquitoes were maintained with a 10% sucrose solution.

### Viral Production and Infection via Bloodmeal

Mayaro virus strain BeAn 343102 (BEI Resources, Manassas, VA, USA) was utilized in this study, a genotype D strain originally isolated from a monkey in Para, Brazil, in May 1978. Virus-infected supernatant was aliquoted and stored at −80°C until used for mosquito infections. Viral stock titers were obtained by focus forming assay (FFA) technique. Adult female mosquitoes at 6 days post emergence that had not previously blood-fed were used for experimentation. Mosquitoes were allowed to feed on either human blood spiked with MAYV at 1*107 FFU/mL or a control bloodmeal with no virus via a glass feeder jacketed with 37°C distilled water for 1 h.

At 2, 7, and 14 days post infection, mosquitoes were anesthetized with triethylamine (Sigma, St. Louis, MO, USA) and RNA was extracted from each individual mosquito using mirVana RNA extraction kit (Life Technologies) applying the protocol for extraction of total RNA. Infection was confirmed via qPCR using primers published by Wiggins et. al. 2018 (Forward: 5′-TGGACCTTTGGCTCTTCTTATC-3′, Reverse: 5′-GACGCTCACTGCGACTAAA-3′) [10], a CT value of 20 or less was used to confirm infection (Supplementary Table 1). 3 pools of total RNA were created for each time point and infection status to be used for library preparation, each consisting of 750 ng of RNA from 4 mosquitoes for a total of 3 mg per pool as confirmed via nanodrop. The protocol for mosquito rearing, viral production, and infection via bloodmeal is described in more detail in Brustolin et al. 2018 [9].

### Transcriptomic Library Preparation and Sequencing

All pools were sent to University of Texas Medical Branch for library preparation where total RNA was quantified using a Qubit fluorescent assay (Thermo Scientific) and RNA quality was assessed using an RNA 6000 chip on an Agilent 2100 Bioanalyzer (Agilent Technologies). See Etebari et all. 2017 for more detail on library preparation and sequencing [17]. 1 mg of total RNA per pool was poly-A selected and fragmented using divalent cations and heat (940 C, 8 min). The NEBNext Ultra II RNA library kit (New England Biolabs) was used for RNA-Seq library construction. Fragmented poly-A RNA samples were converted to cDNA by random primed synthesis using ProtoScript II reverse transcriptase (New England Biolabs). After second strand synthesis, the double-stranded DNAs were treated with T4 DNA polymerase, 5′ phosphorylated and then an adenine residue was added to the 3′ ends of the DNA. Adapters were then ligated to the ends of these target template DNAs. After ligation, the template DNAs were amplified (5-9 cycles) using primers specific to each of the non-complimentary sequences in the adapters. This created a library of DNA templates that have non-homologous 5′ and 3′ ends. A qPCR analysis was performed to determine the template concentration of each library. Reference standards cloned from a HeLa S3 RNA-Seq library were used in the qPCR analysis. Cluster formation was performed using 15.5-17 billion templates per lane using the Illumina cBot v3 system. Sequencing by synthesis, paired end 75 base reads, was performed on an Illumina NextSeq 5500 using a protocol recommended by the manufacturer.

### Small RNA Library Preparation and Sequencing

Small RNA libraries were created using the New England Biolabs small RNA library protocol. See Saldaña et. al. 2017 for more information on small RNA sequencing [21]. Library construction used a two-step ligation process to create templates compatible with Illumina based next generation sequence (NGS) analysis. Where appropriate, RNA samples were quantified using a Qubit fluorometric assay. RNA quality was assessed using a pico-RNA chip on an Agilent 2100 Bioanalyzer. Library creation uses a sequential addition of first a 3′ adapter sequence followed by a 5′ adapter sequence. A cDNA copy was then synthesized using ProtoScript reverse transcriptase and a primer complimentary to a segment of the 3′ adapter. Amplification of the template population was performed in 15 cycles (94°C for 30 sec; 62°C for 30 sec; 70°C for 30 sec) and the amplified templates were PAGE (polyacrylamide gel electrophoresis) purified (147 bp DNA) prior to sequencing. All NGS libraries were indexed. The final concentration of all NGS libraries was determined using a Qubit fluorometric assay and the DNA fragment size of each library was assessed using a DNA 1000 high sensitivity chip and an Agilent 2100 Bioanalyzer. Single end 75 base sequencing by synthesis on an Illumina NextSeq 5500.

### Transcriptomic RNA Sequencing Data Analysis

Raw sequencing data was uploaded to the ICS-ACI high performance computing cluster at Pennsylvania State University to perform all computational analyses. Transcriptomic libraries had adapters trimmed and low-quality bases removed using Trimmomatic read trimming software with base settings [26]. Quality trimmed reads were aligned to the current build of the Anopheles stephensi Indian strain genome in Vectorbase (AsteI2) using the STAR RNA sequencing aligner [27]. Reads less than 75 bp in length and with a mapping quality of less than 20 were dropped from the analysis, and read counts were calculated in R using the rSubread package [28], following which a principal components analysis was performed and differential expression conducted using a negative binomial GLM with the EdgeR package [29]. Contrasts considered in the GLM were infected against control at 2, 7, and 14 dpi, and differences between 2-7 dpi and 7 – 14 dpi for infected treatments corrected for the response from the control treatments between the same time points. Gene IDs that were differentially expressed with a log2FC value of +/− 1 and P value < 0.05 were uploaded to g:Profiler to run GO term overrepresentation analysis [30].

### Small RNA Sequencing Data Analysis

Small RNA libraries had adapters trimmed using Trimmomatic and were subsequently passed into the miRDeep2 pipeline to identify novel and known miRNAs in all samples and determine expression of all known and novel miRNAs at each time point and treatment status [31, 26]. Novel miRNAs with a miRDeep score of less than 3, a minimum free energy value of less than − 20, or a non-significant Randfold p-value were considered false IDs and excluded from further analysis. miRNA targets were identified in the AsteI2 genome using miRanda software package [32]. Differential expression of miRNAs in response to infection status and time point was conducted using a negative binomial GLM with the EdgeR package and contrasts as described for the transcriptomic analysis [29]. miRNAs which were differentially expressed with log2FC +/− 1 and P value < 0.05 had their miRanda genomic targets uploaded to g:Profiler to determine if any GO terms were overrepresented by transcripts potentially regulated by differentially expressed miRNAs [33]. piRNAs and siRNAs were isolated from the small RNA libraries by selecting all 18 – 24 nt reads (siRNA) and 24 – 35 nt (piRNA) reads from the trimmed datasets and filtering out all identified mature miRNAs, and those mapping to the MAYV NC_003417.1 genome were considered potential piRNAs or siRNAs. piRNA and siRNA alignment to the AsteI2 genome was performed using Bowtie RNA sequencing aligner within the MSRG pipeline [34]. Observation of fastQC output for Control Day 7 Replicate 1 small RNA sequencing revealed poor sequencing results, so this replicate was omitted from all analyses in the small RNA focused portions of this study [35].

### Data availability

Sequencing data have been deposited in the GEO depository under accession number GSE165488.

## RESULTS/DISCUSSION

### Transcriptome

#### RNA Sequencing

We assayed genome-wide gene expression in pools of *An. stephensi* (Liston strain) experimentally infected with MAYV at 2, 7, and 14 dpi, along with blood fed uninfected negative controls. RNAseq libraries were sequenced on the Illumina NextSeq 550 platform, yielding 20.6 - 28.4 million paired end reads per library. (Supplementary Table 1). Principal components analysis (PCA) performed on read counts of each annotated gene in the *An. stephensi* (Indian strain) reference transcriptome (AsteI2) at each time point distributed infected and control samples into distinct groups (Figure 1).

**Figure 1:**
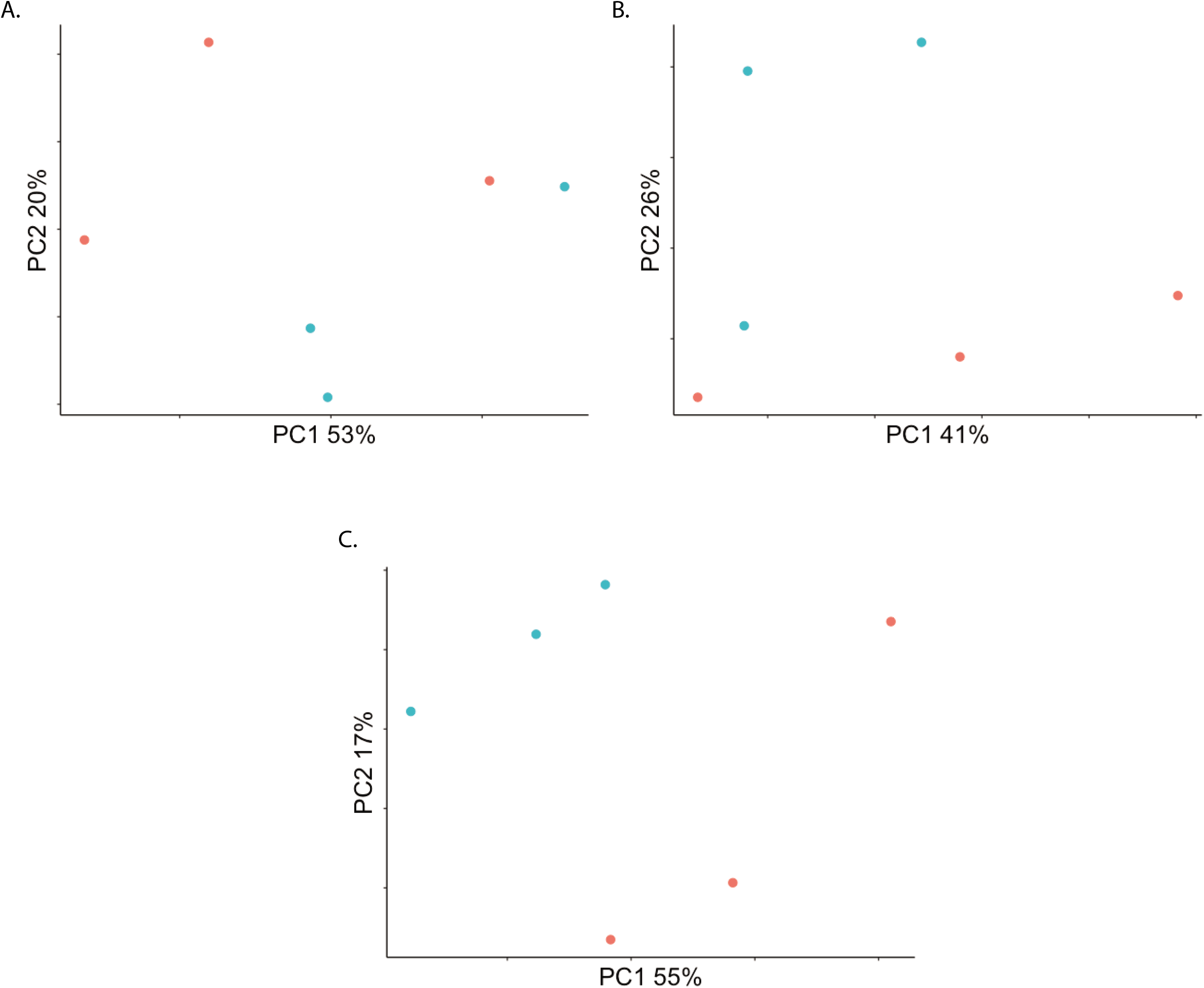
Principal Components Analysis (PCA) on filtered read counts mapping to annotated genes from the AsteI2 build of the *Anopheles stephensi* genome in Vectorbase. A., B., and C. are read counts from samples in the 2, 7, and 14 dpi groupings respectively. In all PCAs, blue is Mayaro infected, and red are control.

#### Differential Expression

To determine which genes exhibit differential expression by infection status and between time points a general-linearized model (GLM) was performed on filtered and normalized read counts mapping to the AsteI2 genome (Supplementary Table 2; Figure 2). Differential expression was also computed for a newer chromosome level genome assembly (Supplementary Table 3), however, the results from the older assembly are discussed here as the genes in the older assembly contain better annotations than those in the newer assembly [36]. Contrasts considered in the GLM were infected compared to control at 2, 7, and 14 dpi, and differences between 2 -7 dpi and 7 – 14 dpi for infected treatments correcting for results from control treatments between those time points. Genes were considered significantly regulated if they had a log fold-change (log2FC) value of +/− 1 and P value < 0.05.

**Figure 2:**
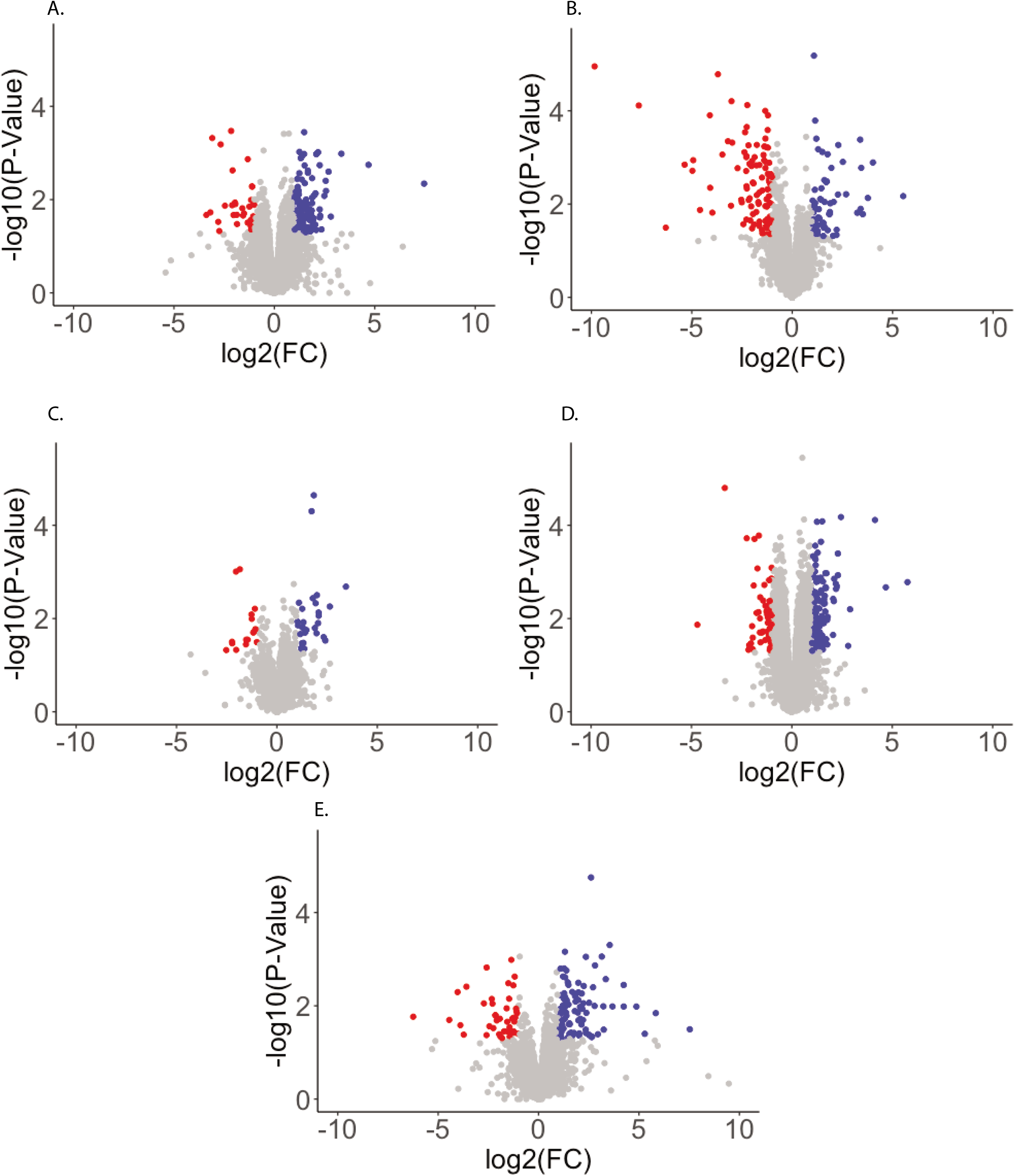
Volcano plots visualizing differential expression of *Anopheles stephensi* transcripts in response to Mayaro infection. The Y-axis shows -log10 transformed P-values, and the X-axis shows log2 transformed fold change values. Red points represent transcripts depleted by more than −1 log2FC in response to infection with a FDR < 0.05, while blue points are transcripts enriched by more than 1 log2FC in response to infection with a P value < 0.05. A. - C. are transcripts regulated in the 2 dpi, 7 dpi, and 14 dpi groupings respectively, while D. and E. are transcripts regulated in the infected treatment between 2 - 7 dpi and 7 - 14 dpi respectively.

There were 161 (64 enriched, 97 depleted), 45 (29 enriched, 16 depleted), and 204 (149 enriched, 55 depleted) of 10,313 annotated genes regulated between control and infected at 2, 7, and 14 dpi respectively. 3 genes were regulated in the same direction at each time point, 2 enriched (ASTEI09037, ASTEI03083) and 1 depleted (ASTEI04716). The gene with the strongest response to infection at any time point was ASTEI04601 at 2 dpi with a log2FC of -9.8 and the most enriched gene was ASTEI04639 at 14 dpi with log2FC of 5.7. When considering changes between time points for the infected treatment when controlling for the response from the uninfected treatments, there were 96 positively and 44 negatively regulated genes between 2 - 7 dpi, and 129 enriched and 32 depleted genes between 7 - 14 dpi. Regulated transcripts for 2-7 dpi ranged from −6.2 (ASTEI08168) to 7.5 (ASTEI09252) log2FC in terms of magnitude of expression, and −3.4 (ASTEI10804) to 7.4 (ASTEI04639) log2FC for 7-14 dpi. When considering a FDR threshold as a multiple testing correction very few transcripts in any contrast can be considered significantly regulated; 3 transcripts at 2 dpi (ASTEI04601, ASTEI05497, ASTEI05732) and 2 transcripts at 14 dpi (ASTEI00644, ASTEI08604) fall below a FDR < 0.1 threshold for significance.

#### Gene Ontology

A gene ontology (GO) over-representation analysis was performed using g:Profiler on gene IDs which were significantly enriched or depleted in any considered contrast in the GLM described above when using a P value cutoff of 0.05 and any overrepresented GO terms with an FDR < 0.5 were considered significant (Supplementary Table 4) [33]. At 2 dpi significantly regulated molecular function terms were overrepresented by peptidase activity, specifically serine type endopeptidase activity, however this overrepresentation is not observed when just considering depleted or enriched genes it is only present when considering all regulated genes at 2 dpi together. Depleted genes at 2 dpi are overrepresented by cell-cell adhesion terms in the biological function category, and components of the membrane for cellular component, enriched genes are not significantly biased for any terms. At 7 dpi molecular function terms related to odorant binding and carboxylic acid binding were overrepresented in depleted genes, and for enriched genes however with a slightly less than significant FDR of 0.06. Molecular function terms relating to dioxygenases, transferases, and oxidoreductases are overrepresented by depleted genes at 7 dpi, and different types of amino acid binding as well as serine type endopeptidase terms are overrepresented by enriched genes. Biological process terms related to nervous system processes, sensory perception of smell, and mannose metabolism are overrepresented by depleted genes at 7 dpi. At 14 dpi the enriched transcripts were biased for molecular function GO terms related to sensory perception, specifically perception of mechanical, light, and sound stimuli, while depleted transcripts were biased for those related to MAPK/JNK signaling cascades, apoptosis, and amino acid biosynthesis for molecular functions and peroxisome and nucleosome for cellular component terms. From 2 to 7 dpi molecular function terms related to serine protease activity were represented by both enriched and depleted genes, but primarily by enriched genes, and enriched genes were biased for cell membrane proteins for cellular component. 7 to 14 dpi had enriched genes biased for molecular functions related to ATP dependent microtubule activity and G-protein coupled receptors while depleted genes were biased for different types of amino acid binding. Biological process terms overrepresented by enriched genes between 7 and 14 dpi were related to sensory perception, and depleted genes were overrepresented for various types of metabolic and biosynthetic processes, NF-kB signaling, and the MAPK/JNK cascade.

Endopeptidases, specifically serine proteases were enriched at 7 dpi and from 2 – 7 dpi, suggesting an activation of the Toll pathway as part of the innate humoral response to infection once the virus has had time to establish an infection in the mosquito [33]. Activation of serine proteases is not uncommon in pathogenic infection of insects and has been identified specifically as enriched in *Ae. aegypti* in response to DENV and Zika virus (ZIKV) infection, and in *An. gambiae* and *An. coluzzii* in response to O’nyong’nyong virus (ONNV) infection [16–18, 37]. At late stages of infection there was depletion of the autophagic and apoptotic inducing JNK and MAPK cascades in addition to repression of JAK/STAT signaling pathways through repression of MAPK signaling. Autophagy and apoptosis both have demonstrated positive impacts on alphaviral replication [33, 38], suggesting another possible molecular response from the mosquito to prevent viral replication at late stages of infection.

### Small RNA

#### miRNA Identification

We next identified novel miRNAs in the small RNA transcriptomes of the MAYV infected samples and controls using miRDeep [31]. We searched for matches in our sequencing reads to all miRNAs in the miRBase database for the species *An. gambiae, Aedes aegypti, Culex quinquefasciatus, Drosophila melanogaster, Bombyx mori, Apis mellifera*, and *Acyrthosiphon pisum*. We found matches to 73 known miRNAs, all from *An. gambiae*, and 78 novel miRNAs identified across all samples, with between 2.2 – 4.0 million reads mapping to identified miRNAs per-sample (Supplementary Table 5 and 6). We found no explicit relationship between diversity of miRNA population and either dpi or infection status. Of the 153 total miRNAs identified across all samples, 83 were present in at least one replicate per treatment (Figure 3). PCA of normalized read counts shows a correlation between infection status and grouping along the PC1/PC2 axis at each time point (Figure 4).

**Figure 3:**
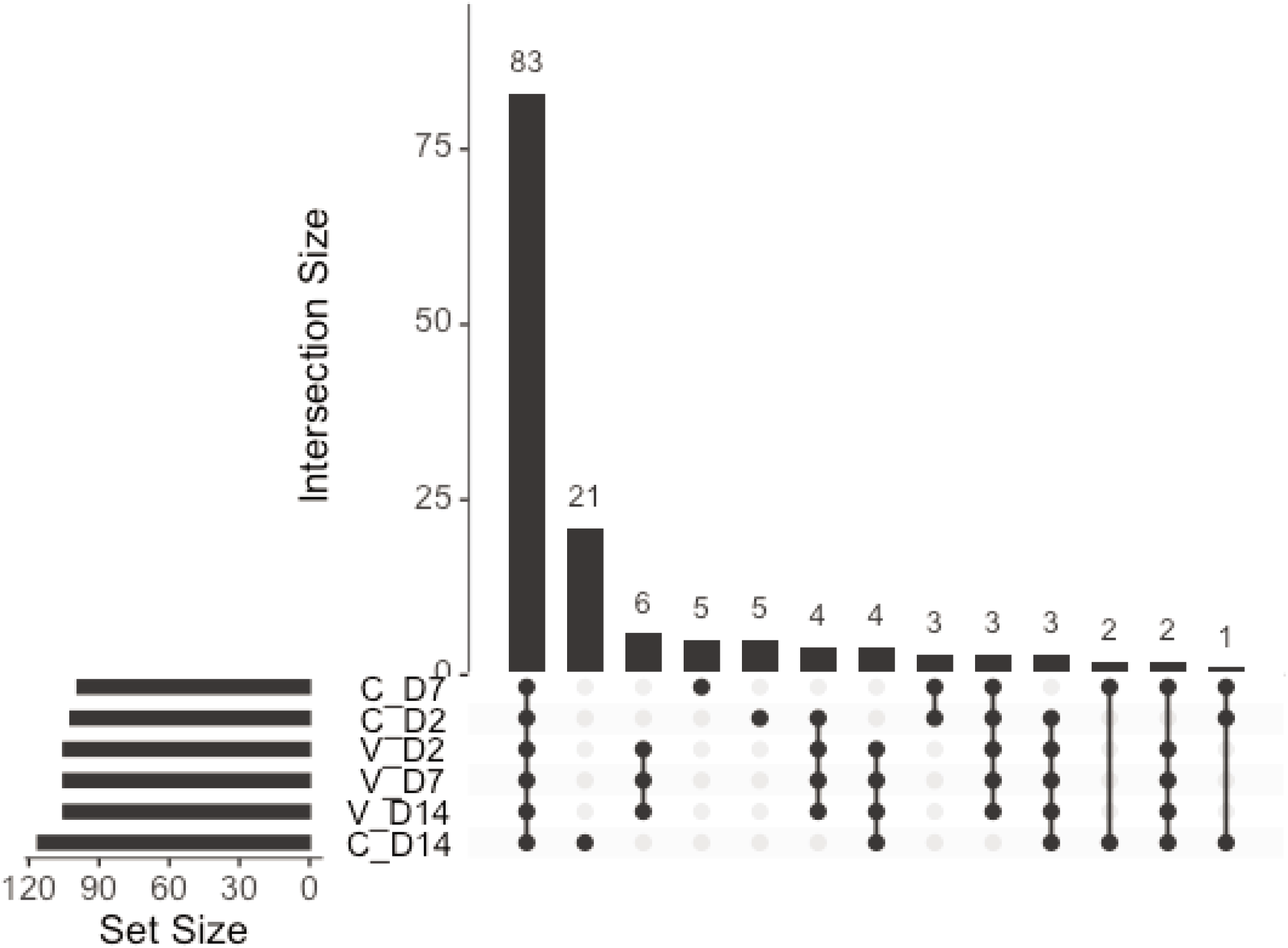
The top histogram represents the number of miRNAs shared between treatments (intersection size), and each row below the histogram represents a treatment. The lines connecting treatments below the top histogram represent treatments which share that number of miRNAs, and the histogram to the side of the treatments represents the number of miRNAs contained within each treatment.

**Figure 4:**
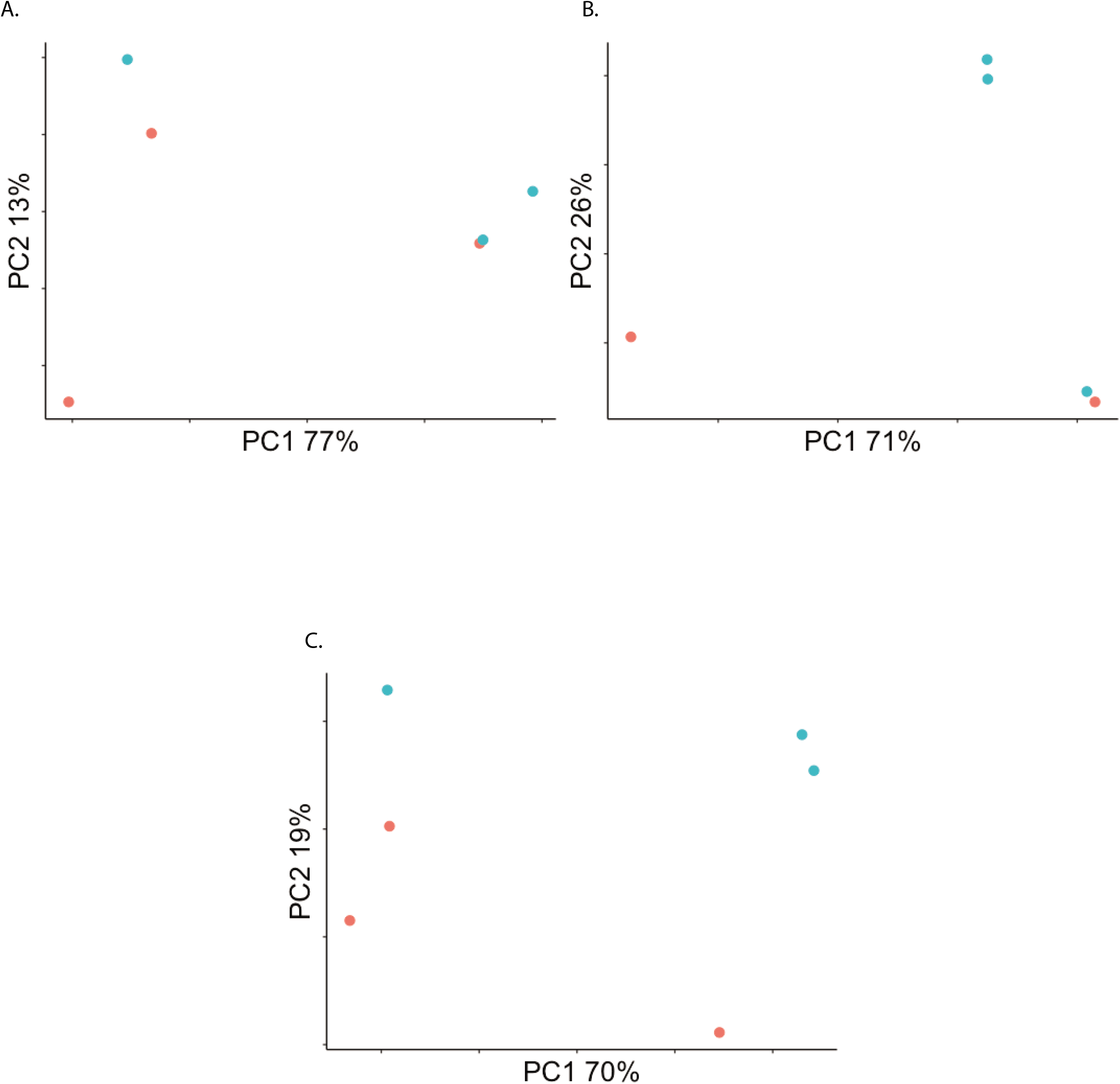
Principal Components Analysis (PCA) on read counts mapping to miRNAs identified in the AsteI2 build of the *Anopheles stephensi* genome in Vectorbase. A. - C. are the 2 dpi, 7 dpi, and 14 dpi groupings respectively. In all PCAs, blue is Mayaro infected, and red is control.

#### miRNA Differential Expression

We next identified known and novel miRNAs that were differentially expressed by infection status (Fig. 5; Table 1, Supplementary Table 7). Contrasts considered in the GLM were infected against control at 2, 7, and 14 dpi, and differences between 2 -7 dpi and 7 – 14 dpi for infected treatments relative to control. miRNAs were considered differentially expressed by having a log fold-change (log2FC) value of +/− 1 and P value < 0.05. There were a total of 8 miRNAs differentially regulated in any considered contrast, novel miRNAs as-mirNOV10, as- mirNOV16, and as-mirNOV17 as well as known miRNAs aga-miR-286b, aga-miR-2944a, aga-miR- 2944b, aga-miR-307, and aga-miR-309. as-mirNOV10 was enriched at 2 dpi, as-mirNOV16 was enriched at 7 dpi, and as-mirNOV 17 was depleted at 14 dpi and between 7 – 14 dpi. The known miRNAs were depleted as a group at 7 dpi and in the 2 - 7 dpi contrast but enriched in the 7 – 14 dpi contrast.

**Figure 5:**
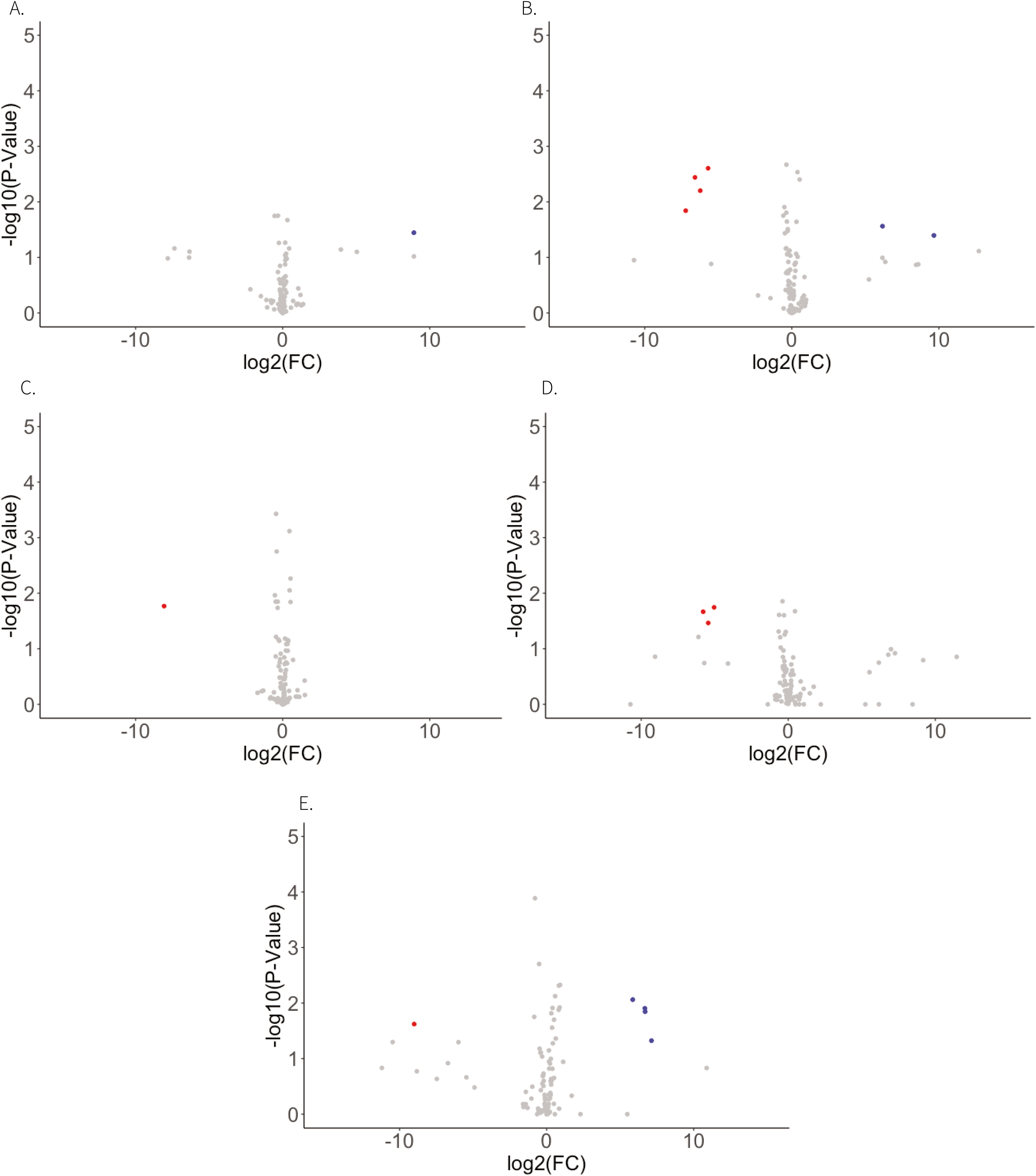
Volcano plots visualizing differential expression of identified *Anopheles stephensi* miRNAs in response to Mayaro infection. The Y-axis shows -log10 transformed P-values, and the X-axis shows log2 transformed fold change values. Red points represent transcripts depleted by more than −1 log2FC in response to infection with a FDR < 0.05, while blue points are transcripts enriched by more than 1 log2FC in response to infection with a FDR < 0.05. A. - C. are the 2 dpi, 7 dpi, and 14 dpi groupings respectively, while D. and E. are miRNAs regulated in the infected treatment between 2 - 7 dpi and 7 - 14 dpi respectively.

**Table 1:** Differentially expressed *Anopheles stephensi* miRNAs in response to Mayaro virus infection.

| miRNA ID | logFC | logCPM | F | P Value | FDR | Direction | Contrast |
| --- | --- | --- | --- | --- | --- | --- | --- |
| as-mirNOV10 | 8.99 | 4.11 | 6.06 | 0.04 | 0.57 | POSITIVE | 2 dpi |
| as-mir1NOV16 | 6.21 | 2.27 | 6.36 | 0.03 | 0.31 | POSITIVE | 7 dpi |
| aga-miR-309 | -7.13 | 10.57 | 16.78 | 0.00 | 0.08 | NEGATIVE | 7 dpi |
| aga-miR-286b | -7.78 | 11.31 | 11.12 | 0.01 | 0.13 | NEGATIVE | 7 dpi |
| aga-miR-2944a | -6.23 | 12.45 | 21.04 | 0.00 | 0.07 | NEGATIVE | 7 dpi |
| aga-miR-2944b | -6.74 | 10.93 | 13.54 | 0.00 | 0.11 | NEGATIVE | 7 dpi |
| as-mirNOV17 | -8.12 | 2.05 | 7.71 | 0.02 | 0.16 | NEGATIVE | 14 dpi |
| aga-miR-286b | -6.67 | 11.31 | 6.10 | 0.03 | 0.52 | NEGATIVE | 2 - 7 dpi |
| aga-miR-2944a | -5.58 | 12.45 | 11.52 | 0.01 | 0.47 | NEGATIVE | 2 - 7 dpi |
| aga-miR-2944b | -5.96 | 10.93 | 7.44 | 0.02 | 0.47 | NEGATIVE | 2 - 7 dpi |
| aga-miR-309 | -6.33 | 10.57 | 9.48 | 0.01 | 0.47 | NEGATIVE | 2 - 7 dpi |
| aga-miR-286b | 7.80 | 11.31 | 7.05 | 0.02 | 0.14 | POSITIVE | 7 - 14 dpi |
| aga-miR-2944a | 6.46 | 12.45 | 14.94 | 0.00 | 0.09 | POSITIVE | 7 - 14 dpi |
| aga-miR-2944b | 7.29 | 10.93 | 10.55 | 0.01 | 0.09 | POSITIVE | 7 - 14 dpi |
| aga-miR-307 | 1.09 | 4.84 | 7.68 | 0.02 | 0.13 | POSITIVE | 7 - 14 dpi |
| aga-miR-309 | 7.30 | 10.57 | 11.72 | 0.01 | 0.09 | POSITIVE | 7 - 14 dpi |
| as-mirNOV17 | -8.76 | 2.05 | 5.42 | 0.05 | 0.21 | NEGATIVE | 7 - 14 dpi |

The miR-309/286/2944 has been found to be enriched in *An. gambiae* in response to blood feeding [39–40], and to be associated with Argonaute proteins post-bloodmeal [39]. When experimentally repressed aga- miR-309 was found to retard oocyte development [39], depletion in response to MAYV infection may suggest that viral replication or the host immune response sequesters resources normally requires for host oocyte development, and as a result associated miRNAs are also depleted. Experimental infections have demonstrated that for *Ae. aegypti*, infection with CHIKV does not impact the number of eggs laid but it does have a detrimental impact on the viability of the eggs produced, however this is not observed with ZIKV [41, 42]. Infection of *Culex tarsalis* with West Nile Virus demonstrates a decrease in fecundity as measured by egg raft size and number of eggs laid [43].

#### miRNA Target Prediction

We next identified putative targets in the *An. stephensi* genome for all known and novel miRNAs identified in all samples [32]. For each of the identified miRNAs we found an average 537 potential annotated targets within the AsteI2 genome (Supplementary Table 8). Targets for significantly regulated miRNAs were loaded into g:Profiler and any overrepresented GO terms with an FDR < 0.5 were considered significant (Supplementary Table 9) [33].

No overrepresented GO terms were identified by the genomic targets of as-mirNOV-10 or aga-miR-307. as-mirNOV-16 was significantly enriched in response to infection at 7 dpi and the only GO terms overrepresented by the predicted genomic targets of this miRNA are associated with protein binding. as-mirNOV17 was depleted at 14 dpi and between 7 – 14 dpi and has GO terms related to transmembrane ion channels overrepresented by its genomic targets. aga-mir-2944a and aga-mir-2944b were both depleted at 7 dpi and between 2 – 7 dpi but enriched between 7 – 14 dpi and both have GO terms primarily associated with intracellular signaling and various binding functions, and aga-mir-2944b also appears to be involved with lipid localization and transport. aga-mir-286b and aga-miR-309 were both also depleted at 7 dpi and between 2 and 7 dpi, and were enriched between 7 and 14 dpi; aga-mir-286b only had acetylglucosaminyltransferase activity overrepresented by its genomic targets, and aga-miR-309 had terms related to calcium ion transport, actin filament binding, and catalytic activity overrepresented by its genomic targets.

The novel miRNA as-mirNOV-17 has 498 predicted genomic targets, and those targets overlap with 8 enriched and 3 depleted genes at 14 dpi and 6 enriched and 2 depleted genes between 7 and 14 dpi, when as-mirNOV-17 was significantly repressed in response to MAYV infection (Table 1). Neither as-mirNOV-10 or as-mirNOV-16 showed a bias for enriched or depleted genomic targets in the contrasts they were significantly regulated in. The known miRNAs also showed a bias for enriched targets between 2-7 dpi and 7-14 dpi where they are repressed and activated in each contrast respectively. These patterns are consistent with the miRNAs acting as effector molecules for RNAi, except for the known miRNAs between 7 - 14 dpi where their expression is enhanced but they still have a bias for enriched genomic targets [38]. Recent studies have demonstrated that through targeting of promotor elements miRNAs can have a positive impact on gene transcription, so this could explain the phenomenon happening between 7 – 14 dpi where miRNA targets are enriched when the miRNAs themselves are also enriched [45].

#### piRNA Identification

We identified putative viral piRNAs in the small RNA datasets by tracking the 24-35 nt reads, removing those that were identified as miRNAs, and mapping these reads to the Mayaro Virus NC_003417.1 genome using the Bowtie sequence aligner, all of which was performed within the MSRG pipeline [26–27, 34, 52]. There was viral piRNA expression in infected samples with a bias for the positive strand over the negative strand, and the proportion of potential piRNAs mapping to the viral genome remained consistent across time points with no particular peaks or hotspots identified across the viral genome (Figure 6) [53].

**Figure 6:**
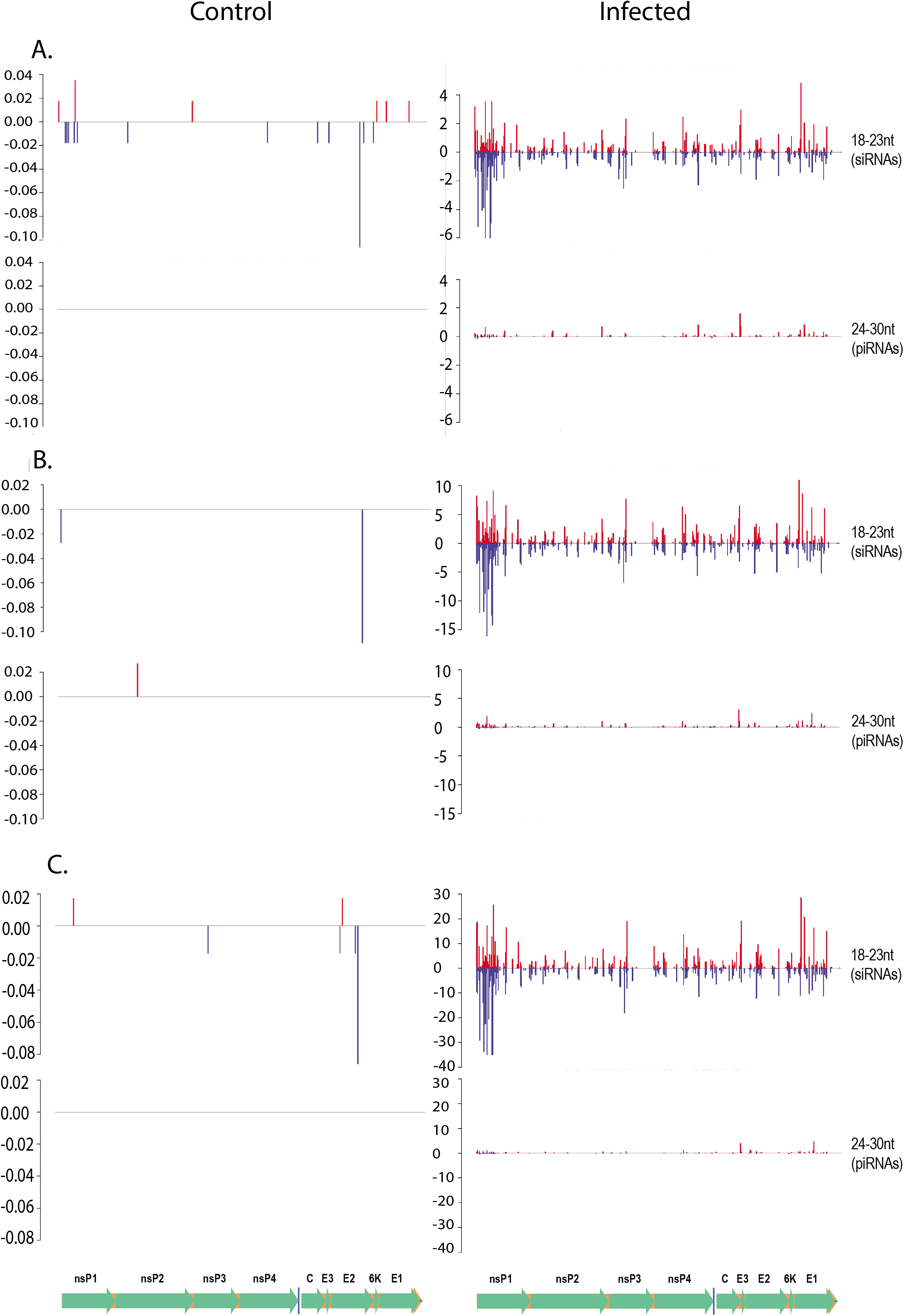
Histograms demonstrating read depth across the Mayaro virus genome for reads with a piRNA size profile (24 - 35 nt) and a siRNA size profile (18 – 23). Y-axis is read depth in reads per million, and X-axis is position in viral genome. Blue demonstrates reads for that sample mapping to the negative strand, while red demonstrates those mapping to the positive strand. A. - C. 2, 7, and 14 dpi respectively, those on the left are control and on the right are infected for that time point. Underneath the y-axis is a visualization of the positions of the proteins along the viral genome.

Virus-derived piRNA-like small RNAs have been identified in insects and insect cells infected with Flaviviruses, Bunyaviruses and Alphaviruses. Knockdown of the piRNA pathway proteins leads to enhanced replication of arboviruses in mosquito cells, suggesting their potential antiviral properties in mosquitoes [46–51]. For example, knockdown of Piwi-4 in *Ae. aegypti* Aag2 cell line increased replication of SFV, and silencing of Ago3 and Piwi-5 led to significantly reduced production of piRNAs against Sindbis virus (SINV) [47,50].

#### siRNA Identification

We also identified putative viral siRNAs in the small RNA datasetsby tracking the 18-24 nt reads that were not miRNAs and could map to the Mayaro Virus NC_003417.1 genome [26, 27, 34, 46]. There was minimal alignment of potential siRNAs to the MAYV genome in the control samples, but in the infected samples there was significant siRNA alignment across the genome (Figure 6). There was a slight negative-strand bias of MAYV siRNAs (Day2 [103+, 124], Day7 [287+, 319−], Day14 [667+, 762−], and the steady increase of siRNAs over time suggests ongoing amplification of the innate immune response with time during the infection process. There were two peaks for siRNA alignment, one at nonstructural protein 1 and another at envelope protein 1 and the number of siRNA reads mapping to the viral genome increased from 2 to 14 dpi.

The siRNA pathway is thought to be the main antiviral component of immunity in insects at the cellular level [54]. Functional studies in *Aedes* mosquitoes have implicated the siRNA pathway as integral to the antiviral response in the midgut stage of infection as well as at later systemic stages of infection for DENV and SINV [54]. Studies of the siRNA response in *An. gambiae* have shown that ONNV does not stimulate a siRNA response at early midgut stages of infection but is present at later stages of infection in the systemic compartments [54]. Our results differ from the findings from *An. gambiae* and ONNV in that the siRNA response is detectable at early stages of infection and is persistent until later stages.

### Conclusion

The transcriptomic profiles suggest that MAYV activates the Toll pathway at early and mid-stages of infection as an innate humoral response from the host to fight infection. At later stages of infection there appears to be a repression of JNK and MAPK signaling cascades, potentially impacting autophagic and apoptotic processes as a way to limit MAYV replication. The small RNA profiles suggests a reliance on siRNA silencing as an antiviral immune response. There was a minimal piRNA response to infection with a positive strand bias, but it did not appear to play a major role in antiviral immunity in this study. miRNAs were also elicited in response to infection and some overlap was observed with transcripts identified as regulated in response to infection, but not to the extent that they appear to be strongly regulating transcriptional profiles in response to infection.

## Acknowledgments

We thank the UTMB sequencing core for assistance with next generation sequencing. This work was supported by NIH grants R01AI150251, R01AI128201, R01AI116636, USDA Hatch funds (Project #PEN04769), and a grant with the Pennsylvania Department of Health using Tobacco Settlement Funds to JLR, NIH grant R01GM135215 to NCL, BBSRC awards BB/T001240/1 and BB/V011278/1, Royal Society Wolfson Fellowship RSWF\R1\180013, NIH grants R21AI138074, URKI grants 20197 and 85336, EPSRC grant V043911/1, and NIHR grant NIHR2000907 to GLH. CAH was supported by an NSF graduate research fellowship program award (ID 2018258101). SH was supported the Liverpool School of Tropical Medicine Director’s Catalyst Fund award. GLH is affiliated to the National Institute for Health Research Health Protection Research Unit (NIHR HPRU) in Emerging and Zoonotic Infections at University of Liverpool in partnership with Public Health England (PHE), in collaboration with Liverpool School of Tropical Medicine and the University of Oxford. GLH is based at LSTM. The views expressed are those of the author(s) and not necessarily those of the NHS, the NIHR, the Department of Health or Public Health England. We would also like to thank Dr Martin Donnelly at Liverpool School of Tropical Medicine for comments on an early version of this manuscript which helped us to identify an error in the analysis.

---

Supplementary Table 1: Information related to infection of *Anopheles stephensi* with Mayaro virus. Includes number of mosquitoes in each treatment and time point and associated mortality, nanodrop readings for all RNA extractions collected, pooling scheme for sequencing of mRNA and small RNA, and qPCR data from each sample using primers specific for Mayaro virus strain BeAn to confirm infection status.

Supplementary Table 2: Differentially expressed transcripts from the *Anopheles stephensi* AsteI2 genome.

Supplementary Table 3: Differential expression results from the *Anopheles stephensi* PRJNA629843 genome.

Supplementary Table 4: GO term overrepresentation for differentially regulated transcripts.

Supplementary Table 5: Read counts mapping to the identified *Anopheles stephensi* miRNAs in each small RNA sample sequenced, both raw counts and counts normalized to library size (reads per million, RPM) are presented.

Supplementary Table 6: All known and novel mature and precursor miRNA sequences identified.

Supplementary Table 7: Differential expression of *Anopheles stephensi* miRNAs.

Supplementary Table 8: Genomic targets from the *Anopheles stephensi* AsteI2 genome for all identified miRNAs.

Supplementary Table 9: Overrepresented GO terms represented by targets of significantly regulated miRNAs.

